# PHYTOMap: Multiplexed single-cell 3D spatial gene expression analysis in plant tissue

**DOI:** 10.1101/2022.07.28.501915

**Authors:** Tatsuya Nobori, Marina Oliva, Ryan Lister, Joseph R. Ecker

## Abstract

Retrieving the complex responses of individual cells in the native three-dimensional tissue context is crucial for a complete understanding of tissue functions. Here, we present PHYTOMap (Plant HYbridization-based Targeted Observation of gene expression Map), a multiplexed fluorescence *in situ* hybridization method that enables single-cell and spatial analysis of gene expression in whole-mount plant tissue in a transgene-free manner and at low cost. We applied PHYTOMap to simultaneously analyze 28 cell type marker genes in *Arabidopsis* roots and successfully identified major cell types, demonstrating that our method can substantially accelerate the spatial mapping of marker genes defined in single-cell RNA-seq datasets in complex plant tissue.

## Main

Understanding how individual cells respond and interact with each other in the face of changing environments is the cornerstone of understanding tissue function. Single-cell transcriptomics technologies have been widely adopted in plant research, enabling the classification of cells into populations that share molecular features for the in-depth analysis of cell types and states^1-3^. The increasing throughput and sensitivity in single-cell transcriptomics technologies will offer tremendous granularity at which cells can be classified, but it will also create new challenges in dealing with cell populations that our current histological and physiological understanding of plant cells cannot account for. To understand the identity and function of molecularly defined cell populations, it is critical to analyze their spatial localization in the tissue.

In plant research, the most common tool for spatially mapping cell population marker genes identified in single-cell transcriptome analysis has been transgenic reporter lines that express fluorescent proteins under the predicted promoter region of the genes; in most cases, each transgenic line visualizes the expression of only one gene. This approach has several limitations in analyzing cells in complex tissue: (1) A cell type/state is not always defined by the expression of a single gene but by the combination of many genes. (2) Spatial mapping of a single gene or a few genes has difficulties in analyzing multiple cell types/states simultaneously, which is critical for understanding interactions between cell types/states. (3) The generation of transgenic plants is time-consuming. (4) The heterologous expression of fluorescent proteins does not necessarily reflect the true expression of the gene because the reporter cassettes lack the native genomic context (e.g., enhancer-promoter interactions). Therefore, spatial gene expression analysis needs to be done with a large number of genes at single-cell resolution for a more complete understanding of the function of cell types/states and their interactions with other cells and the environment.

Spatial transcriptomics technologies hold great promise in addressing these problems by simultaneously revealing the molecular details and spatial location of cells in complex tissues. Methods employing spatially barcoded arrays or imaging-based highly multiplexed smFISH (single-molecule fluorescence *in situ* hybridization) allow researchers to study the expression of many genes (from dozens to the whole transcriptome) with spatial information (from tissue region to single-cell levels)^4^, and such technologies have recently been adopted in plant research^5-7^. Although spatial transcriptomics, combined with single-cell transcriptomics, will contribute to elucidating the spatial organization of cell types/states in plants in great detail^8^, tissue types amenable for spatial transcriptomics experiments are limited to thin (single-cell layer) sections, posing challenges in its application in plants and other organisms. For instance, the root tip—an important organ for plant growth, nutrient acquisition, and interactions with microbes—is a difficult tissue for sectioning due to its small size. Moreover, sectioning will lead to the loss of information from the other parts of the tissue, which may contain the cell types/states of interest; information about environments, such as microbial colonization, can also be lost by sectioning. It may be possible to overcome these problems by sampling serial sections and conducting multiple experiments followed by the 3D reconstitution of 2D data, but spatial transcriptomics technologies are highly costly, making such an approach rarely affordable. To overcome these limitations, we introduce PHYTOMap (Plant HYbridization-based Targeted Observation of gene expression Map), a low-cost single-cell spatial gene expression analysis that can simultaneously map dozens of genes in whole-mount plant tissue.

PHYTOMap builds upon *in situ* hybridization techniques in plants^9,10^ and *in situ* sequencing technologies primarily developed in neuroscience fields^11^. After fixing whole-mount plant tissues, DNA probes (SNAIL probes) with gene-specific barcodes are specifically hybridized on target mRNA molecules, circularized, and amplified *in situ* (Fig. 1a, Methods for details). The hybridization condition has been optimized to allow high target specificity (Fig. S1b). The amplification of DNA barcodes provides high signal-to-noise ratio, enabling signal detection from cleared whole-mount tissue. The location of mRNA molecules is defined by the sequence-by-hybridization (SBH) chemistry^12^ that targets the barcode sequences of DNA amplicons across sequential rounds of probing, imaging, and stripping (Fig. 1b). In each imaging round, four targets are detected using each of the four channels of a confocal microscope (Supplementary Movie 1). After imaging, fluorescent detection probes are stripped (Fig. S1c), and the next round of hybridization targets a new set of four genes (Fig. 1b and 1c). A previous study that used the SBH chemistry to detect amplified DNA probes *in situ* showed that signal was maintained at least over ten sycles^12^.

**Fig. 1.**
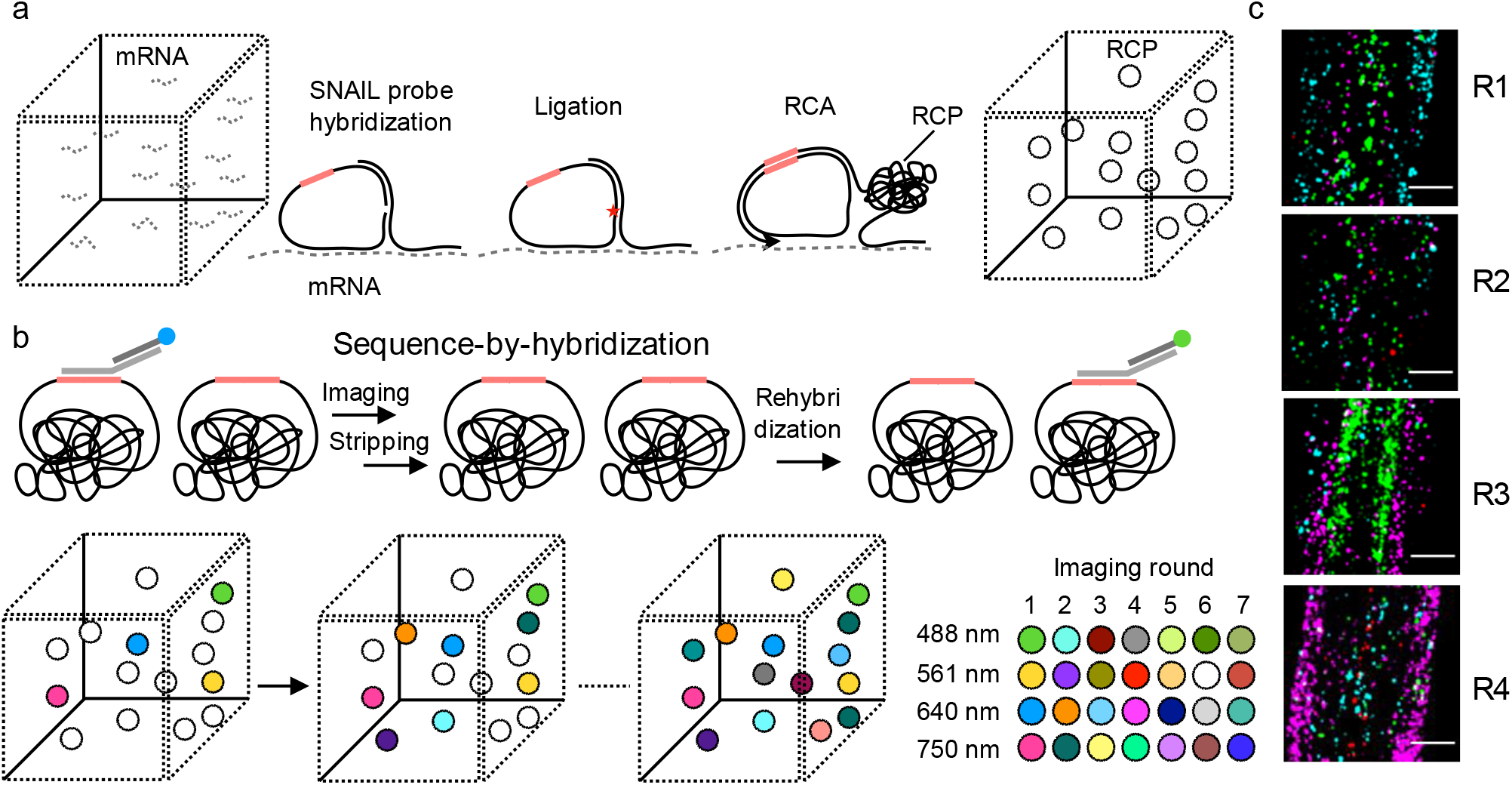
PHYTOMap principles. a, In fixed whole-mount tissue, target mRNA molecules are hybridized by pairs of DNA probes (SNAIL probes) that harbor mRNA species-specific barcode sequences (pink bars). The barcode-containing DNA probes are circularized by ligation (a red star) and amplified *in situ* by rolling circle amplification (RCA). During amplification, amine-modified nucleotides are incorporated into the DNA amplicons (rolling cycle products; RCPs) and stably cross-linked with the cellular protein matrix using a nonreversible amine cross-linker. Amplified DNA barcodes are detected by sequence-by-hybridization chemistry (see b) through multiple rounds of imaging. b, Sequence-by-hybridization. Before each imaging round, four kinds of bridge probes are hybridized to a set of four DNA barcodes. Then, each bridge probe is targeted by one of four fluorescent probes to be imaged. After the imaging, bridge probes and fluorescent probes are stripped away, while keeping RCPs in place. These steps are repeated until all the DNA barcodes are read. c, Representative images at different imaging rounds. Maximum exposure of 60 z-planes of the same position in tissue is displayed. Scale bars = 30 µm.

PHYTOMap successfully mapped well-established/validated cell type marker genes in expected cell types/regions in the root tip of *Arabidopsis* (Fig. 2a-f). The marker genes we targeted include AT4G28100 (EN7; Endodermis) AT4G29100 (BHLH68; Pericycle), AT5G37800 (RSL1; Trichoblast), AT5G53730 (NHL26; Xylem), AT5G57620 (MYB36; Endodermis), AT5G58010 (LRL3; Trichoblast), and AT3G54220 (SCR; Endodermis) (Fig. 2b-e)^13,14^. PHYTOMap also validated cell type/region marker candidates predicted in a previous scRNA-seq study of *Arabidopsis* root tips^15^. For instance, AT3G46280 was detected in the root cap and elongating epidermis as predicted in the scRNA-seq data (Fig. 2d). Genes enriched in meristematic (AT5G42630) and elongation (AT5G12050) zones in the scRNA-seq data were mapped in the expected regions (Fig. 2e); AT5G12050 signal was detected in epidermis and vasculature as shown in scRNA-seq (Fig. 2e). QC and columella signal was also detected from the marker genes AT2G28900, AT3G20840, and AT3G55550 (Fig. 2f). Taken together, PHYTOMap can be used as an efficient tool for validating marker genes identified in scRNA-seq data without generating transgenic plants.

**Fig. 2.**
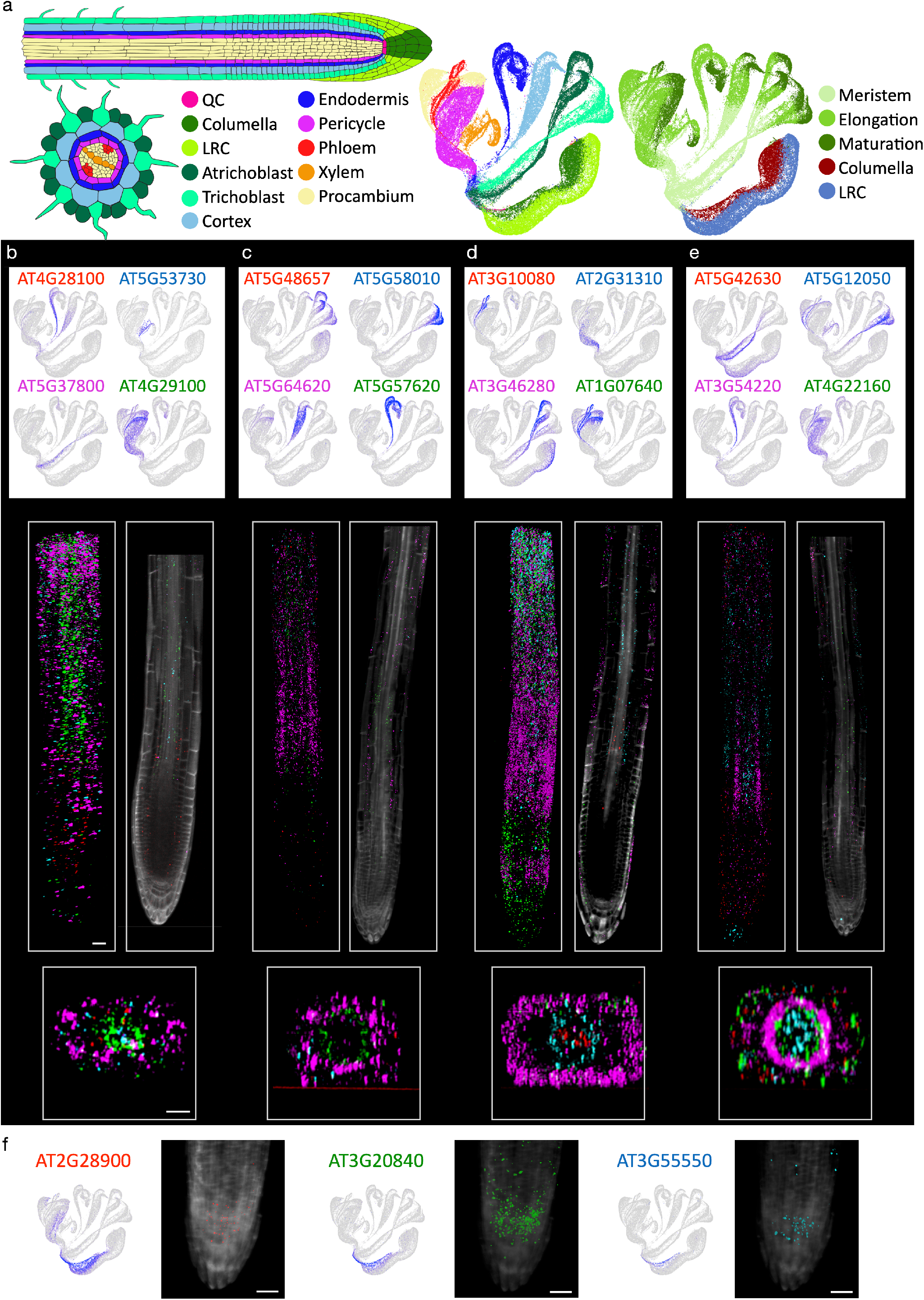
Whole-mount spatial mapping of root tip cell type marker genes predicted in scRNA-seq data with PHYTOMap. a, Schematic representation of the root tip and UMAPs displaying root tip scRNA-seq data used in this study. In the UMAPs, cells are labeled with cell types (left) and regions (right). QC: Quiescent Center. LRC: Lateral Root Cap. b-e, Representative results from each imaging round. Top: UMAPs showing expression patterns of target genes. The colors of gene name labels correspond to the colors in the images below. Middle: 3D projections (left) and optical sections (2D, right) of whole-mount tissue images. Bottom: Representative cross-section views of the middle part of the samples (transition/elongation zone). f, Validated and predicted marker genes for QC and columella. 3D images were shown with cell wall staining. Scale bars = 25 µm.

To demonstrate the multiplexing capacity of this method, we simultaneously targeted 28 genes in the same root tips with seven rounds of imaging. The targeted genes include known cell type marker genes as well as unvalidated cell type marker candidates identified in the scRNA-seq data^15^ (see a full list in Table 1), which showed varying levels of expression in the root tip (Fig. S2). We developed a computational pipeline to integrate whole-mount images from each imaging round and analyze gene expression at the single-cell resolution (Fig. 3a, Methods for details). Cell wall boundary information was obtained together with the RNA-derived signal in each imaging round to facilitate this process. The analysis pipeline first registers 3D images across imaging rounds using cell boundary information, automatically detects spots derived from single mRNA molecules, and annotates spots with gene names. A merged image with detected and decoded transcripts successfully captured the cell type architecture of the root tip (Fig. 3b). To analyze the spatial data at the single-cell level, cell segmentation was performed based on cell wall boundary information using PlantSeg, which performs deep learning-assisted cell boundary prediction and graph partitioning-based cell segmentation^16^ (Fig. 3a). Annotated spots were assigned to individual cells and counted, resulting in a cell-by-gene matrix, a standard scRNA-seq data form that can be used for clustering and dimension reduction analyses (Fig. 3a).

**Table 1:** List of genes analyzed in this study.

| gene ID | gene name | Cell type |
| --- | --- | --- |
| AT2G28900 | OEP161 | Meristematic-Columella |
| AT3G20840 | PLT1 | QC |
| AT1G71692 | AGL12 | Elongating-Phloem |
| AT2G34140 | CDF 4 | Stele/procambium etc |
| AT3G10080 | AT3G10080 | Elongating-Xylem |
| AT1G07640 | OBP2 | phloem |
| AT3G46280 | AT3G46280 | Elongating-Atrichoblast |
| AT2G31310 | LBD14 | pericycle |
| AT5G42630 | KAN4 | Meristematic-Atrichoblast |
| AT4G22160 | AT4G22160 | Elongating-Pericycle |
| AT3G54220 | SCR | Meristematic-Endodermis |
| AT5G12050 | AT5G12050 | Elongating-Trichoblast |
| AT5G48657 | AT5G48657 | Mature-Atrichoblast |
| AT5G57620 | MYB36 | Elongating-Endodermis |
| AT5G64620 | C/VIF2 | Elongating-Cortex |
| AT5G58010 | BHLH82 | Mature-Trichoblast |
| AT4G30080 | ARF16 | Root cap |
| AT1G07710 |  | phloem |
| AT1G16390 | OCT3 | differentiating endodermis |
| AT1G71930 | VND7 | Xylem |
| AT1G79840 | GL2 | Atrichoblast |
| AT2G40260 |  | Cortex |
| AT2G46570 | LAC6 | Xylem |
| AT3G55550 | LECRK-S.4 | Quiescent Center + Columella |
| AT4G28100 | EN7 | Endodermis |
| AT4G29100 | BHLH68 | Pericycle |
| AT5G37800 | RSL1 | Trichoblast |
| AT5G53730 | NHL26 | Xylem |

**Fig. 3.**
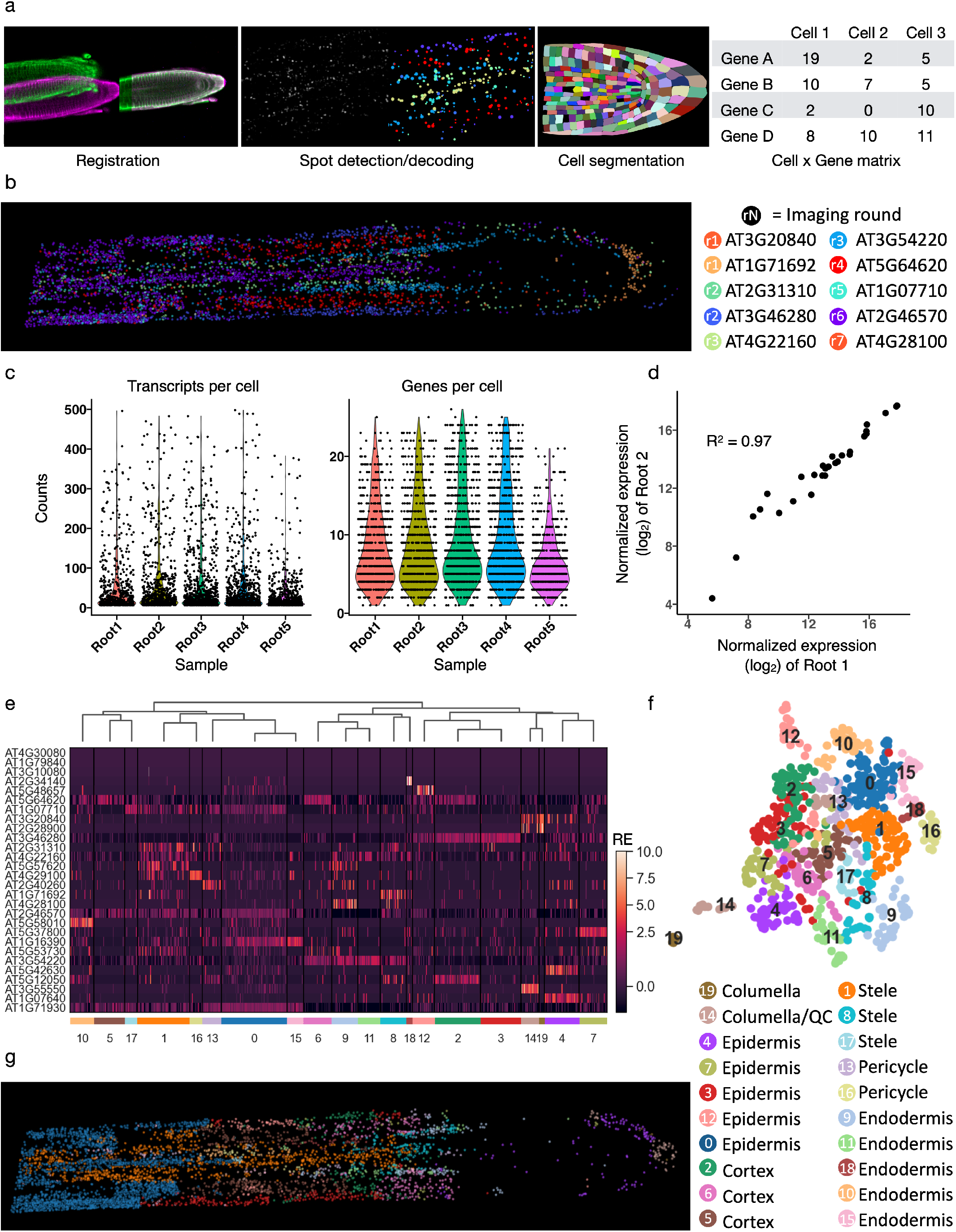
Single-cell and spatial analysis of 28 genes with PHYTOMap. a, PHYTOMap data analysis pipeline for single-cell analysis. b, 3D visualization of transcripts detected and decoded after image registration in a representative root tip (Root 4). A middle section (90-120th z-planes out of 208 z-planes) of the image is displayed. Representative genes from each imaging round were shown. c, Violin plots showing the number of unique RNA molecules (left) and genes (right) detected in five root tip samples. d, Scatter plot comparing normalized bulk expression of each gene between two samples (Root 1 and Root 2). e, Hierarchical clustering of 3608 cells from five root tips based on the relative expression (RE) of 28 genes. Cluster IDs are indicated at the bottom. f, UMAP visualization of the clusters shown in (e). g, 3D visualization of transcripts colored by clusters in (e, f) in a representative root tip (Root 4). A middle section (90-120th z-planes out of 208 z-planes) of the image is displayed.

We analyzed five root tip preparations and identified a total of 259,781 RNA molecules from 3608 cells (median 19 molecules per cell) (Fig. 3c). The assays were highly robust and reproducible, detecting comparable numbers of transcripts for each RNA species between different biological samples (Fig. 3d). This suggests that gene expression between cells or samples can be compared quantitatively. Hierarchical clustering and heatmap visualization revealed cell population-specific expression of target genes (Fig. 3e). We performed de novo clustering using PHYTOMap data and visualized the data on UMAP without using any spatial information (Fig. 3f). These clusters successfully captured major cell types and developmental stages in the root tip (Fig. 3g). Together, these results demonstrate that PHYTOMap can spatially map dozens of genes at the single-cell resolution in a highly reproducible manner.

In conclusion, PHYTOMap is a novel technology that enables multiplexed single-cell spatial gene expression analysis in whole-mount plant tissue without requiring transgenic plant lines. A PHYTOMap experiment can be performed on a timescale similar to other *in situ* hybridization protocols in *Arabidopsis* ^9,10^; sample preparation takes 4-5 days with ∼10 h total bench time (Fig. S3a). Imaging can be performed with a regular confocal microscope. Each imaging round takes 3 h for one root tip and 5 h for five root tips in the present study; thus 21 h and 35 h to finish imaging for a 28-gene experiment in one and five root tips, respectively. Previous studies have shown that more than 25 rounds of imaging are possible with DNA amplicons obtained with approaches similar to our method ^17^, suggesting that the current version of PHYTOMap can target more than 100 genes. The cost of PHYTOMap experiments is approximately $80 for a 28-gene experiment and $230 for a 96-gene experiment (Fig. S3b), where each experiment can accommodate five or more root tips, which can be from different treatments and/or genotypes. Importantly, the transgene-free nature of PHYTOMap makes this technology potentially applicable to any plant species. Cell type annotation in scRNA-seq is challenging in many crop plants as their marker genes are often not conserved in other well-characterized species such as *Arabidopsis*. We believe that PHYTOMap will become a widely used tool for efficient cluster annotation in scRNA-seq studies of a variety of plant species. Beyond cell typing, PHYTOMap will offer unique opportunities to interrogate spatial regulation of complex cellular responses in plant tissue during stress and development.

## Methods

### Sample preparation

#### Cell type marker gene analysis

The *Arabidopsis thaliana* accession Col-0 seeds (hereafter Arabidopsis) were sown on square plates containing Linsmaier and Skoog (LS) medium (Caisson, LSP03) with 0.8% sucrose solidified with 1% agar (Caisson Labs, A038). Plates were kept vertically for five days in a growth chamber under an 8-hour light/16-hour dark regime at 21°C.

### PHYTOMap experimental procedure

#### Chemicals and enzymes

Poly-D-Lysine coated dish (MatTek, P35GC-1.5-14-C). T4 DNA Ligase (Thermo Scientific, EL0011). EquiPhi29 DNA Polymerase (Thermo Scientific, A39391). SUPERaseIn RNase Inhibitor (Invitrogen, AM2696). Aminoallyl dUTP (AnaSpec, AS-83203). Dulbecco’s Phosphate Buffered Saline (DPBS) (Sigma, D8662). BSA, molecular biology grade (New England Biolabs, B9000S). dNTPs (New England Biolabs, N0447S). Fluorescent Brightener 28 disodium salt solution (Sigma, 910090). Paraformaldehyde (Sigma, F8775). Triton-X (Sigma, 93443). Proteinase K (Invitrogen, 25530049). Nuclease-Free Water (Invitrogen, AM9937). BS(PEG)9 (Thermo Scientific, 21582). 20 ×SSC buffer (Sigma-Aldrich, S6639). Ribonucleoside vanadyl complex (New England Biolabs, S1402S). Formamide (Sigma, F9037). Tris, pH 8.0, RNase-free (Invitrogen, AM9855G). EDTA, pH 8.0, RNase-free (Invitrogen, AM9260G). Cellulase (Yaklut, YAKL0013). Macerozyme (Yakult, YAKL0021). Pectinase (Fisher Scientific, ICN19897901).

#### Targeting probe design

Target genes were manually selected based on their cell type-specific expression. Probes were constructed by combining the probe design used in STARmap^18^ and HYBISS^12^ (Fig. S1a). A SNAIL probe is a pair of a padlock probe (PLP) and a primer designed as follows: (1) For each gene, 40-50 nt sequences with a GC content of 40-60% were selected and confirmed that there is no homologous region in the other transcripts by blasting against TAIR10 Arabidopsis genome; (2) selected sequences were split into halves of 20-25 nt (the 5’ halves for PLPs and the 3’ halves for primers), with 2 nt gap in between, and ensured that the melting temperature (*T*_*m*_) of each half is around 60°C; (3) PLPs have complementary sequences for target specific bridge-probes; (4) for each gene, four SNAIL probes were designed; (5) PLPs and primers have complementary sequences to form a circular structure. Bridge probes and detection read-out probes were designed as described before^12^ and detailed in Table 1. All probes were manufactured by Integrated DNA Technologies (IDT).

#### Sample fixation and permeabilization

Five-day-old root tips were mounted on a Poly-D-Lysine coated dish, and all the following steps were conducted on the dish. Samples were fixed, dehydrated, and rehydrated in a similar way as described in previous studies^19,20^ with modifications. Arabidopsis root tips were immersed in FAA (16% (v/v) formaldehyde, 5% (v/v) acetic acid, 50% ethanol) for 1 hour at RT. RNase-free water was used throughout the entire protocol. Samples were then dehydrated in a series of 10 min washes once in 70% (v/v in Nuclease-free water) ethanol, once in 90% ethanol, and twice in 100% ethanol, followed by 10 min washing twice in 100% methanol and stored in 100% methanol at -20°C overnight. On the next day, samples were rehydrated in a series of 5 min washes in 75% (v/v), 50%, and 25% methanol in DPBS-T (0.1% Tween 20 in DPBS) at RT. The cell wall was partially digested by incubating samples in cell wall digestion solution (0.06% cellulase, 0.06% macerozyme, 0.1% pectinase, and 1% SUPERase in DPBS-T) for 5 min on ice, then 30 min at RT. After 2x washes in DPBS-TR (DPBS-T and 1% SUPERase), samples were fixed in 10% (v/v) formaldehyde for 30 min at RT and washed with DPBS-TR. Proteins were digested by incubating samples in Protein digestion buffer (0.1M Tris-HCl (pH 8), 50mM EDTA (pH 8)) with 1:100 volume of Proteinase K (20mg/ml, RNA grade, Invitrogen, 25530049) for 30 min at 37°C. After 2x washes in DPBS-TR, samples were fixed in 10% (v/v) formaldehyde for 30 min at RT and washed with DPBS-TR.

#### SNAIL probe hybridization, amplification, and fixation

The following steps are based on STARmap protocols^18^ with modifications. A pool of SNAIL probes (500 nM each) was heated at 90°C for 5 min and cooled down at RT. Samples were incubated in hybridization buffer (2X SSC, 30% formamide, 1% Triton-X, 20 mM Ribonucleoside vanadyl complex, and pooled SNAIL probes at 10 nM per oligo) in a 40°C humidified oven overnight. After hybridization, samples were washed twice in DPBS-TR and once in 4X SSC in DPBS-TR for 30 min at 37°C and rinsed with DPBS-TR at RT. Samples were then incubated in a T4 DNA ligation mixture (1:50 dilution of T4 DNA ligase supplemented with 1X BSA and 0.2 U/µL of SUPERase-In) at RT overnight. After ligation, samples were washed twice with DPBS-TR for 10 min at RT and incubated in a rolling circle amplification (RCA) mixture (1:20 dilution of equiPhi29 DNA polymerase, 250 µM dNTP, 0.1 µg/ µL BSA, 1 mM DTT, 0.2 U/µL of SUPERase-In, and 20 µM Aminoallyl dUTP) at 37°C overnight. After RCA, samples were rinsed in DPBS-T and covalently crosslinked with 4.3 µg/µL BS(PEG)9 in DPBS-T. BS(PEG)9 was then quenched by incubating samples in 1 M Tris-HCl (pH 8) for 30 min at RT.

#### Gel embedding and tissue clearing

After the fixation of DNA amplicons, samples were embedded in acrylamide gel by incubating in a polymerization mixture (4% Acrylamide, 0.2% bis-acrylamide, 0.1% ammonium persulfate, and 0.1 % tetramethylethylenediamine in DPBS-T) for 1.5 hours at RT. Samples were then rinsed in DPBS-T. After gel embedding, samples were cleared by incubating in ClearSee^21^ at RT overnight.

#### Sequence-by-hybridization

Samples were washed with 2X SSC for 5 min at RT and then incubated in a bridge probe hybridization mixture (2X SSC, 20% formamide, and four bridge probes at 100 nM per oligo in water) for 1 hour at RT. After washing twice in 2X SSC for 5 min at RT, samples were incubated in a detection probe hybridization mixture (2X SSC, 20% formamide, 1:100 dilution of Calcofluor White (Fluorescent Brightener 28 disodium salt solution), and fluorescent detection oligos at 100 nM per oligos in water) for 1 hour at RT. Samples were washed in 2X SSC and ClearSee for 5 min at RT and stored in ClearSee until imaging.

#### Imaging

Imaging was performed using Leica Stellaris 8 confocal microscope equipped with DMi8 CS Premium, Supercontinuum White Light Laser, Laser 405 DMOD, Power HyD detectors, and an HC PL APO CS2 ×40/1.10 water objective. Image pixel size was 512×1536 px with voxel size of 0.57 µm x 0.57 µm x 0.42 µm. The following channel settings were used: 405 nm Ex, 420-510 nm EM; 499 nm Ex, 504-554 nm EM; 554 nm Ex, 559-650 nm EM; 649 nm Ex, 657-735 nm EM; 752 nm Ex, 760-839 nm EM.

### PHYTOMap data processing

#### Image registration

Sample handling could cause shifts in a field-of-view (FOV) during image acquisition. To correct these shifts, image stacks from each round were registered in 3D based on the cell wall boundary staining information by a global affine alignment using random sample consensus (RANSAC)-based feature matching ^22^. We adopted the analysis pipeline of *Bigstream*^23^ with modifications. The first round of images was used as a reference. The registered images were used for downstream analysis with *starfish* (https://github.com/spacetx/starfish), a Python library for processing image-based spatial transcriptomics data.

#### Spot detection and decoding

Images were denoised using the *Bandpass* function, and the z-axis was smoothed by Gaussian blurring using the *GaussianLowPass* function. Using the *Clip* function, an image clipping filter was applied to remove pixels with too low or too high intensity. FISH signals (spots) from single molecule-derived rolling circle products were detected by a blob detection technique using the *BlobDetector* function, which is a multi-dimensional gaussian spot detector that convolves kernels of multiple defined sizes with images to identify spots. The kernel sizes were determined based on the diameter of spots (typically around 1 µm). Detected spots were decoded based on the imaging round and the channel information using the *SimpleLookupDecoder* function.

#### Cell segmentation

The cell wall staining image of the first imaging round (the same image used as a reference for image registration) was used for segmentation. The PlantSeg workflow^16^ was used to predict cell boundaries and label the cells in the image stacks. The re-scaling factor of [1.68, 2.28, 2.28] was used to fit our images to the “confocal_PNAS_3d” model on the software. Graphics processing unit (GPU)-based convolutional neural network (CNN) prediction was used for cell boundary prediction with the patch size of [80, 160, 160] and the “accurate” mode (50% overlap between patches). The MultiCut segmentation algorithm with Under/Over-segmentation factor=0.5, 3D watershed, CNN Predictions Threshold=0.3, Watershed Seeds Sigma=1.0, Watershed Boundary Sigma=0, Superpixels Minimum Size=1, Cell Minimum Size=1. After segmentation, images were re-scaled with the appropriate factors.

#### Spot assignment to segmented cells

Based on segmentation masks generated in the previous step, individual decoded spots were assigned to cells by using the *AssignTargets* function. Then, spots were counted for each target in each cell, resulting in a cell-by-gene matrix.

#### Image visualization

Registered and decoded images were visualized with *napari*^24^, a fast, interactive, multi-dimensional image viewer for Python, by using the *starfish* function *display*.

### PHYTOMap count data analysis

*scanpy* was used for analyzing count data^25^. Cells that contain fewer than six spots (transcripts) were filtered out from the analysis. Count data were log-transformed, and principal components (PCs) were calculated. A neighborhood graph was computed by using 10 PCs with a local neighborhood size of five. UMAP embedding was generated based on the neighborhood graph. Clustering was performed with the Leiden algorithm with the parameter resolution=1.

### scRNA-seq analysis

Processed and annotated data by Shahan et al.^15^ was downloaded from GEO (GSE152766_Root_Atlas_spliced_unspliced_raw_counts.rds.gz). The R package Seurat (v4.1.0)^26^ was used to display the expression of target genes.

### Code availability

The code to analyze PHYTOMap data is available at https://github.com/tnobori/PHYTOMap.

## Acknowledgments

We thank Joanne Chory and Xuelin Wu (Salk) for letting us use their confocal microscope, Robert Henley (Salk) for useful discussions, and Ecker lab members for the critical comments on the manuscript. T.N. was supported by Human Frontiers Science Program (HFSP) Long-term Fellowship (LT000661/2020-L). J.R.E. is an Investigator of the Howard Hughes Medical Institute.

## Author contributions

T.N. conceived and designed the study and experiments with guidance from J.R.E.; T.N. performed experiments, analyzed data, and wrote the manuscript; M.O. and R.L. optimized tissue preparation protocols; M.O., R.L., and J.R.E edited the manuscript.

---

**Extended Data Fig. 1.**
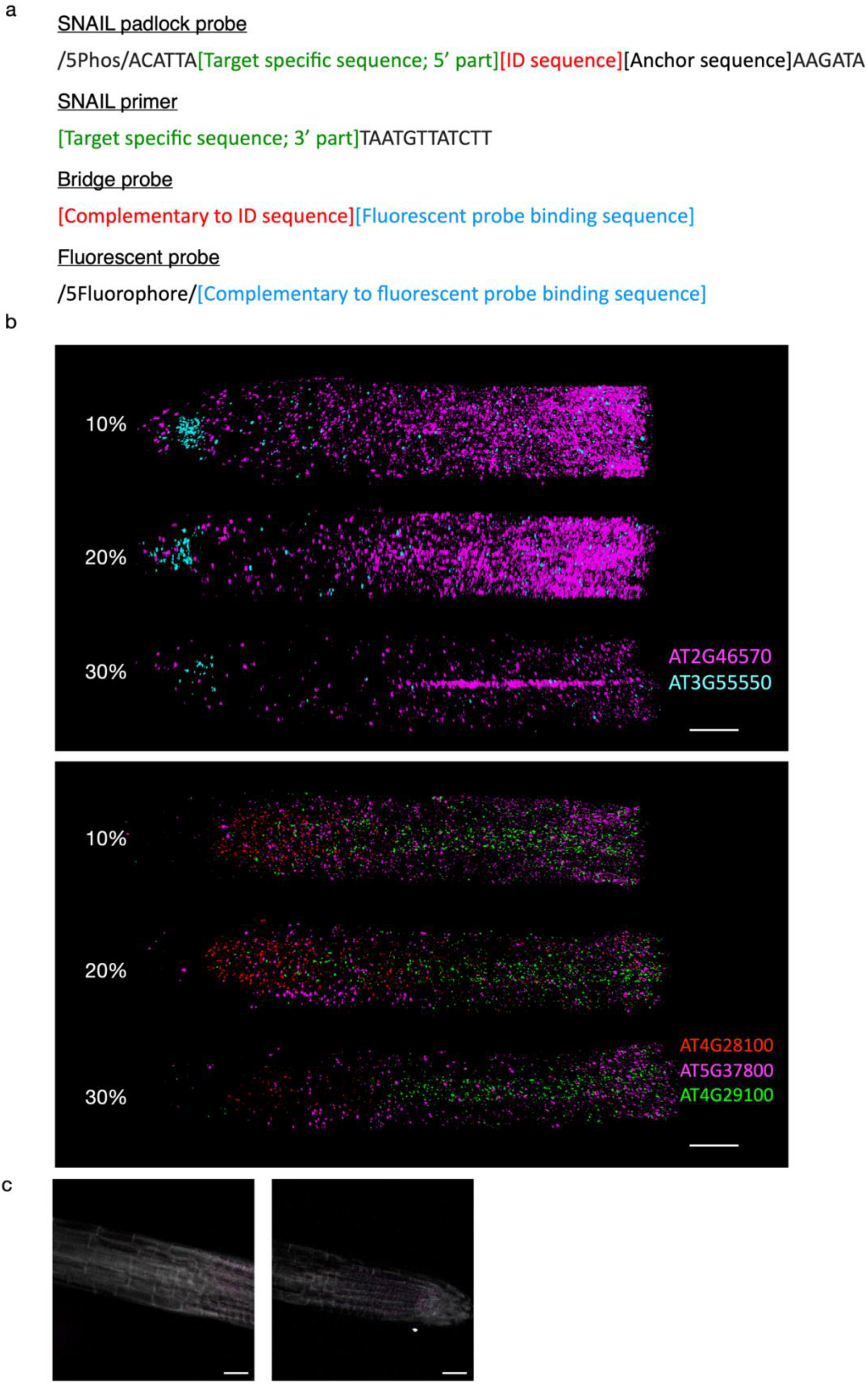
PHYTOMap method development. a, Design of probes used in PHYTOMap. ID sequences are unique to different RNA species. Anchor sequence was included based on a previous study^12^ but not used in the present study. b, Optimization of formamide concentration during SNAIL probe hybridization. Hybridization in 30% formamide showed higher target specificity. c, Images after stripping fluorescent probes. The four-color channels are shown in higher contrast than in Fig. 2b, and cell wall staining images are overlaid. Scale bars = 25 µm.

**Extended Data Fig. 2.**
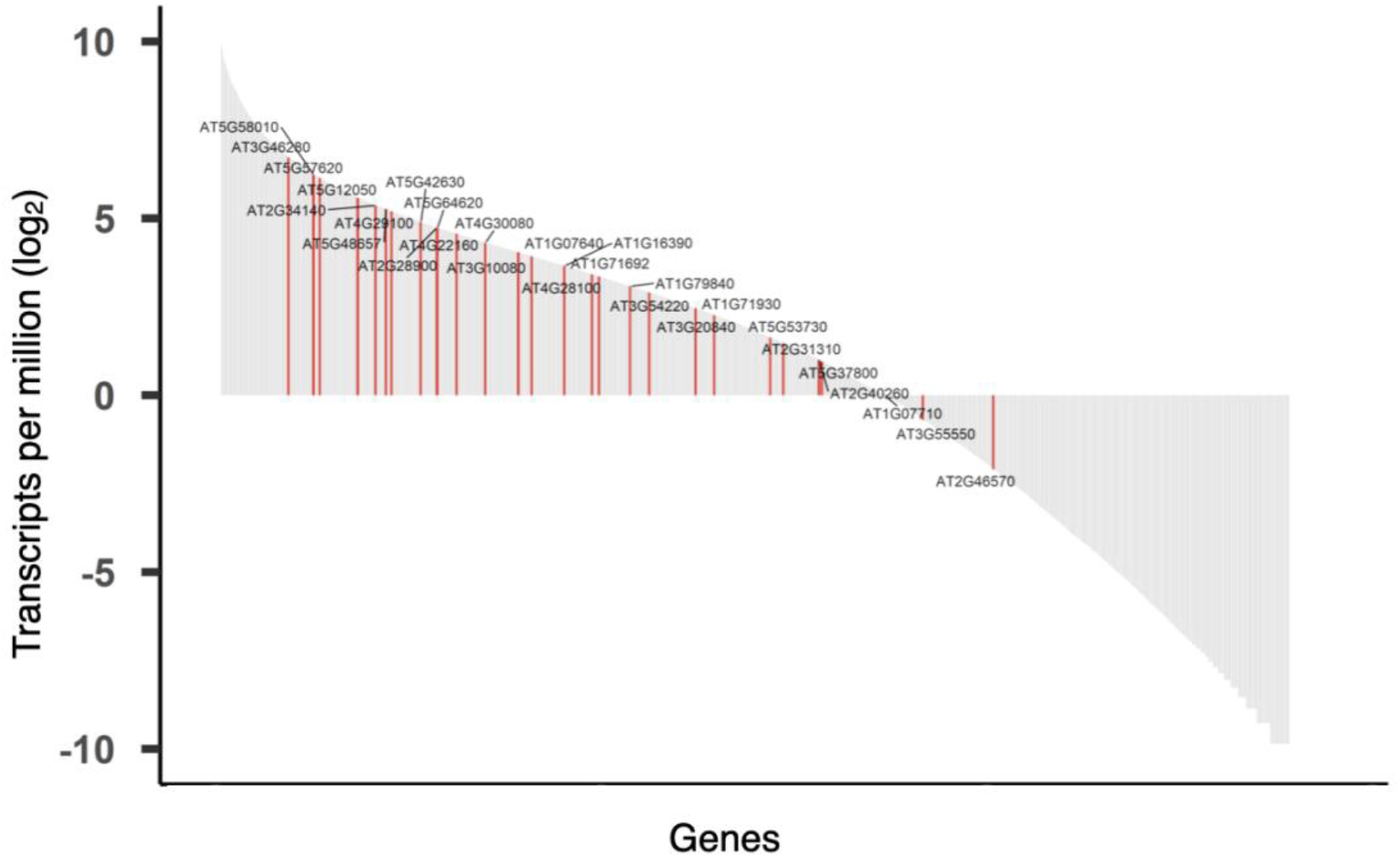
Varying levels of expression of the genes targeted in this study. Bulk expression levels of genes (transcript per million) were calculated based on the root tip scRNA-seq data^15^. The twenty-eight genes targeted in this study were highlighted in red.

**Extended Data Fig. 3.**
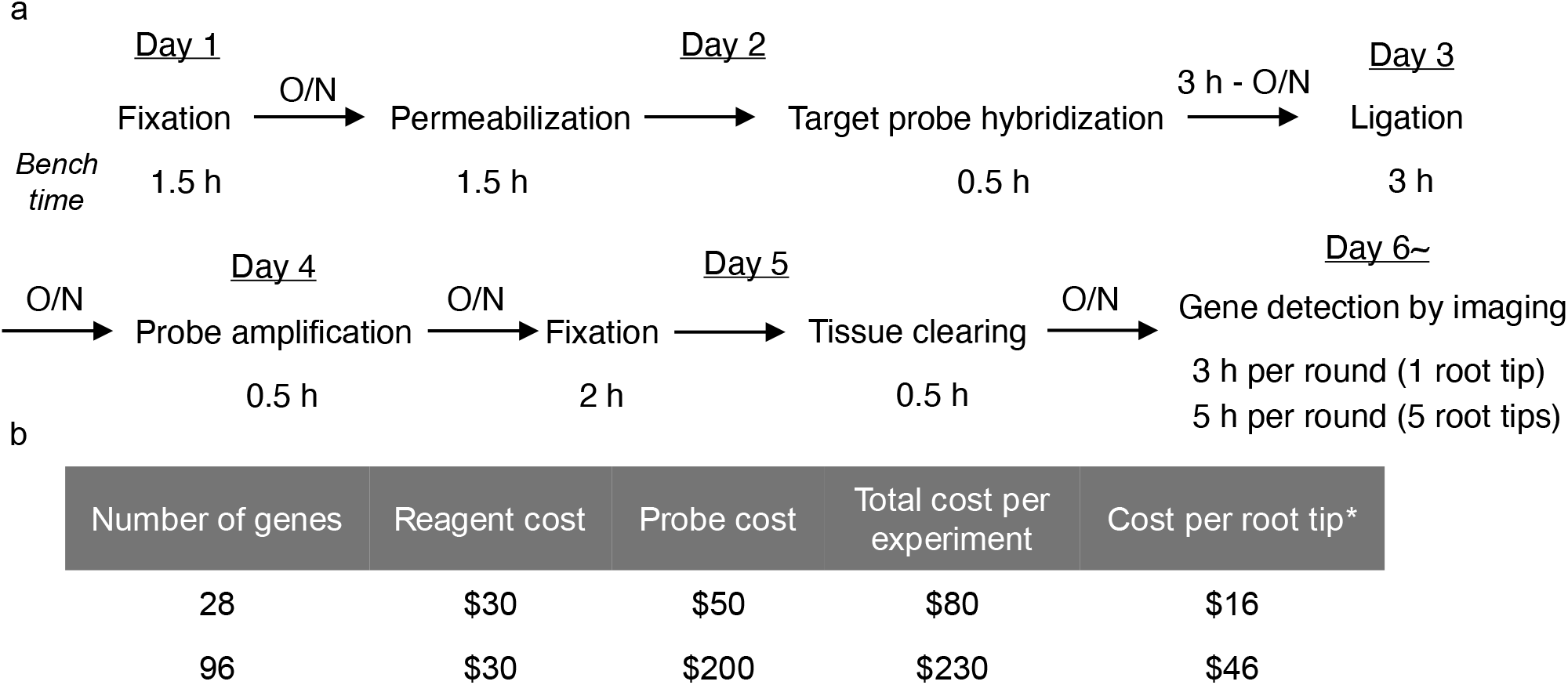
Timing and cost estimates of PHYTOMap. a, Diagram showing the key steps and timings of PHYTOMap. b, Cost estimates for PHYTOMap experiments at different scales. A sample that fits within a 14 mm diameter circle can be analyzed in one experiment using the protocols presented in this study. *Calculated for five root tips per experiment. It is possible to accommodate more root tips in one experiment.

**Supplementary Table** | **DNA probes used in this study**

**Supplementary Movie** | **PHYTOMap 3D images**

Representative 3D images of root tips visualized by PHYTOMap.

